# MARSY: A multitask deep learning framework for prediction of drug combination synergy scores

**DOI:** 10.1101/2022.06.07.495155

**Authors:** Mohamed Reda El Khili, Safyan Aman Memon, Amin Emad

## Abstract

**Motivation:** Combination therapies have emerged as a treatment strategy for cancers to reduce the probability of drug resistance and to improve outcome. Large databases curating the results of many drug screening studies on preclinical cancer cell lines have been developed, capturing the synergistic and antagonistic effects of combination of drugs in different cell lines. However, due to the high cost of drug screening experiments and the sheer size of possible drug combinations, these databases are quite sparse. This necessitates the development of transductive computational models to accurately impute these missing values.

**Results:** Here, we developed MARSY, a deep learning multi-task model that incorporates information on gene expression profile of cancer cell lines, as well as the differential expression signature induced by each drug to predict drug-pair synergy scores. By utilizing two encoders to capture the interplay between the drug-pairs, as well as the drug-pairs and cell lines, and by adding auxiliary tasks in the predictor, MARSY learns latent embeddings that improve the prediction performance compared to state-of-the-art and traditional machine learning models. Using MARSY, we then predicted the synergy scores of 133,722 new drug-pair cell line combinations, which we have made available to the community as part of this study. Moreover, we validated various insights obtained from these novel predictions using independent studies, confirming the ability of MARSY in making accurate novel predictions.

**Availability and Implementation:** An implementation of the algorithms in Python and cleaned input datasets are provided in https://github.com/Emad-COMBINE-lab/MARSY.

**Supplementary Information:** Online-only supplementary data is available at the journal’s website.

## Introduction

Cancers are complex diseases that involve various pathways and are regulated by a multitude of different genes (Li, et al., 2020; Sun, et al., 2015). Despite the emerging understanding of cancers, the development of effective treatments remains a prevailing challenge. Combination therapies, in which multiple treatments are administered simultaneously, have emerged as an alternative to monotherapies (Bayat Mokhtari, et al., 2017; Madani Tonekaboni, et al., 2018; Sun, et al., 2015). By simultaneously targeting multiple genes, proteins, and pathways, they can reduce the probability of drug resistance while improving treatment’s efficacy (Bayat Mokhtari, et al., 2017; Li, et al., 2018; Li, et al., 2018; Li, et al., 2021), and allow for the reduction of necessary dosage per drug, reducing the risk of drug toxicity and adverse effects (Kuru, et al., 2022; Li, et al., 2018; Li, et al., 2021).

Due to the distinct molecular and clinical characteristics of cancer types, it is necessary to evaluate the response of cancer cells to different treatments and treatment strategies in each cancer type. The curation of molecular profiles of cancer cell lines (CCLs) and their response to monotherapies in large databases such as the Cancer Cell Line Encyclopedia (CCLE) (Barretina, et al., 2012) and Genomics of Drug Sensitivity in Cancer (GDSC) (Yang, et al., 2013) initiated the development of various computational models for prediction of single drug response in CCLs (Costello, et al., 2014; Hostallero, et al., 2022) and patient tumours (Hostallero, et al., 2021; Huang, et al., 2020). More recently, large databases of synergy scores of drug combinations (mainly drug-pairs) in CCLs such as DrugComb (Zagidullin, et al., 2019) have been curated based on results of many high-throughput drug screening studies. These databases include different synergy scores to quantify the observed effectiveness of a drug combination compared to its expected effectiveness if they are not interacting (Amzallag, et al., 2019). In spite of these efforts, the sheer size of possible (drug combination, CCL) tuples and the cost of drug screening experiments have resulted in sparse datasets with many missing values. Computational models that can accurately predict the synergy scores of drug combinations and impute these datasets can have a significant impact in this domain.

Various models have been proposed to achieve this goal (Bansal, et al., 2014). In one study (Sidorov, et al., 2019), the authors used various machine learning (ML) models and obtained their best predictive performance using Random Forests (RF) and Extreme Gradient Boosting (XGB) models. Other studies have also explored ML models such as linear regression, Support Vector Machine (SVM), LASSO, RF, XGB, Extremely Randomized Trees, and TreeCombo (which is an extreme gradient boosted tree-based approach) (Celebi, et al., 2019; Janizek, et al., 2018; Jeon, et al., 2018). In addition to these traditional ML models, several deep learning (DL) models have been proposed. One of the earliest methods, DeepSynergy (Preuer, et al., 2017), used a fully connected neural network composed of two hidden layers and showed significant improvement compared to then state-of-the-art ML models such as RF and XGB. A more recent model, MatchMaker (Kuru, et al., 2022), showed that learning two distinct representations for each drug-CCL pair in a combination can improve the performance.

Here we present MARSY, a multitask DL model that predicts the level of synergism between drug-pairs tested on CCLs. Using gene expression to characterize CCLs and drug-induced signatures to represent each drug, MARSY learns a distinct set of embeddings to obtain multiple views of the input features. Precisely, a representation of the entire combination and a representation of the drug-pair are learned in parallel. These embeddings are then fed to a multitask network that predicts the synergy score of the drug combination alongside single drug responses. A thorough evaluation of MARSY revealed its superior performance compared to various state-of-the-art and traditional computational methods. A detailed analysis of the design choices of our framework demonstrated the predictive contribution of the learned embeddings by this model. Using MARSY, we then predicted the synergy scores of 133,722 new drug-pair CCL combinations, which can be used to guide future drug screening and pharmacogenomics studies. Moreover, we validated various insights obtained from these novel predictions using independent studies, confirming the ability of MARSY in making accurate novel predictions.

## METHODS

### Prediction of drug-pair synergy scores and single drug response using MARSY

MARSY (<u>M</u>ultitask drug p<u>A</u>i<u>R</u> <u>Sy</u>nergy) is a DL-based model that seeks to learn latent representations (embeddings) capable of predicting the synergy score of two drugs in a CCL. As input, it receives feature vectors of two candidate drugs and a feature vector of a CCL (Figure 1). The architecture includes two parallel and separately parameterized encoders with a bottleneck layer, and one multi-task predictor. The first encoder (ENC_Pair_) receives the concatenation of the feature vectors of the drug-pair to learn a (drug 1, drug2)-specific embedding, while the second encoder (ENC_Triple_) receives all three feature vectors to learn a (drug1, drug2, CCL)-specific embedding. These embeddings are then concatenated and provided as input to the predictor, which predicts the synergy score of the two drugs, along with the single drug response of each drug in that CCL (performing three tasks simultaneously). The inclusion of single drug response predictors as auxiliary tasks ensures that the representations learned by the two encoders are constrained to capture biologically and chemically important information corresponding to each drug, improving its generalizability and performance.

**Figure 1:**
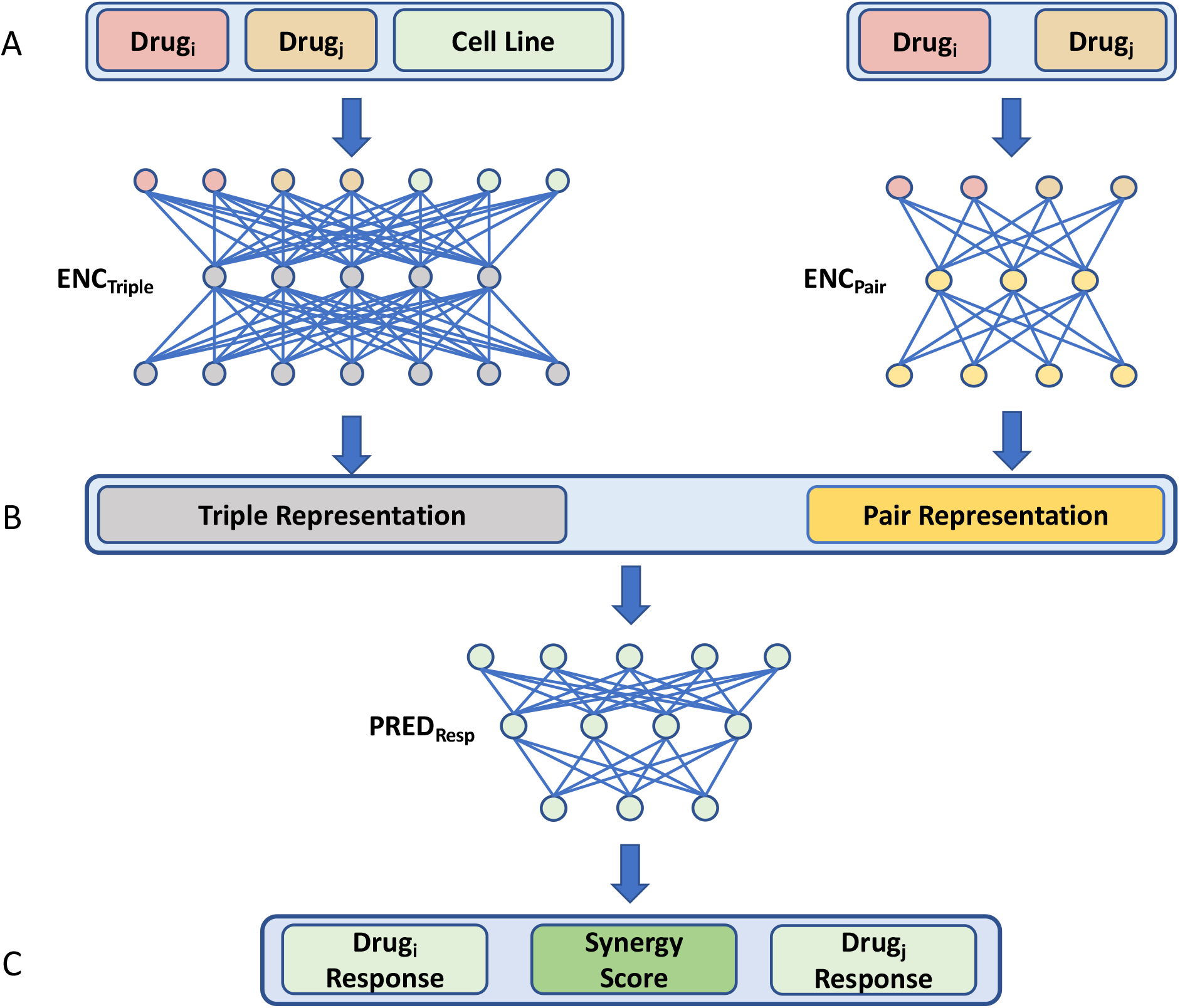
The overview of MARSY’s architecture. A) The drug features and CCL gene expression profile are provided as input to the two encoders. Separately parameterized encoders obtain representations corresponding to the drug-pair and to the drugs and CCL triple. B) Both representations are concatenated and used as inputs to a multi-task predictor. C) The predictor predicts the synergy score of the drug combination, along with the single drug response of each drug (three outputs).

ENC_Pair_ accepts a drug-pair’s signature (a vector of length 3912) as input and consists of two fully connected hidden layers of width 1024 and 2048. ENC_Triple_ accepts a vector of length 8551 corresponding to the concatenation of a drug-pair’s signatures and a CCL’s gene expression profile as input and consists of two fully connected hidden layers of width 2048 and 4096. The first hidden layer of both encoders is a bottleneck layer to perform dimensionality reduction, while the second hidden layer has a larger width to increase the capacity of the model. The relative width of layers in these two encoders are chosen to approximately match the length of inputs to each encoder. A linear activation function is used for the first layer, while a Rectified Linear Unit (ReLU) function is used for the second layer of both encoders. Both encoders use dropout regularization with a probability of 0.2. The embeddings obtained from these two encoders are concatenated and provided as input to the predictor (PRED_Resp_). PRED_Resp_ consists of two fully connected hidden layers of width 4096 and 1024 and an output layer of width 3 to predict the synergy score of the drugs and their individual drug response. A ReLU activation function is used for the first two layers while a linear activation function is selected for the output layer, given the regression nature of the prediction task. Finally, a dropout regularization with a probability of 0.5 is used for the predictor. Details of hyperparameter tuning and training are discussed below.

### Training and hyperparameter tuning

Both encoders (ENC_Triple_ and ENC_Pair_) and the predictor (PRED_Resp_) were trained in an end-to-end fashion using a mean squared error (MSE) loss function and the Adamax optimizer (Kingma and Ba, 2014) with the default learning rate of 0.001 and early stopping. The dropout probabilities and the activation function used on the input layer of both encoders were considered hyperparameters to be tuned using an independent validation set. All possible combinations of the probabilities 0.2 and 0.5 were assessed for both the encoders and the predictor. Furthermore, this assessment was performed alternating between a ReLU and a linear activation function for the input layer of the encoders. The reason we considered a linear activation function as an option for the first layer is its ability to improve and simplify the optimization process. These hyperparameters were evaluated on a small validation set (^~^1100 samples) that was excluded from the main dataset used for cross-validation evaluation, in order to ensure that no data leakage occurs. The rest of MARSY’s architecture followed the design choices detailed above. The best performing combination of hyperparameters (as evaluated on the independent validation set) was used to finalize the design of the MARSY framework.

### Datasets and data cleaning

In this study, the feature vector of each CCL corresponded to its baseline gene expression profile (before administration of any drug), as measured in CCLE (Barretina, et al., 2012). We downloaded the *log*_2_(*RPKM* + 1) normalized RNA-seq profile of 1019 untreated CCLs (catalogued in the CCLE study) from the CellMiner database (Reinhold, et al., 2012; Shankavaram, et al., 2009). We removed lowly expressed genes whose *log*_2_(*RPKM* + 1) value was lower than 1 in more than 75% of all CCLs. Additionally, we calculated the variance of genes across all CCLs and excluded those that had a variance smaller than 0.8, which resulted in a set of 4639 features (Figure 2). Finally, a z-score normalization was performed to the expression of each gene over all the CCLs, one gene at a time.

**Figure 2:**
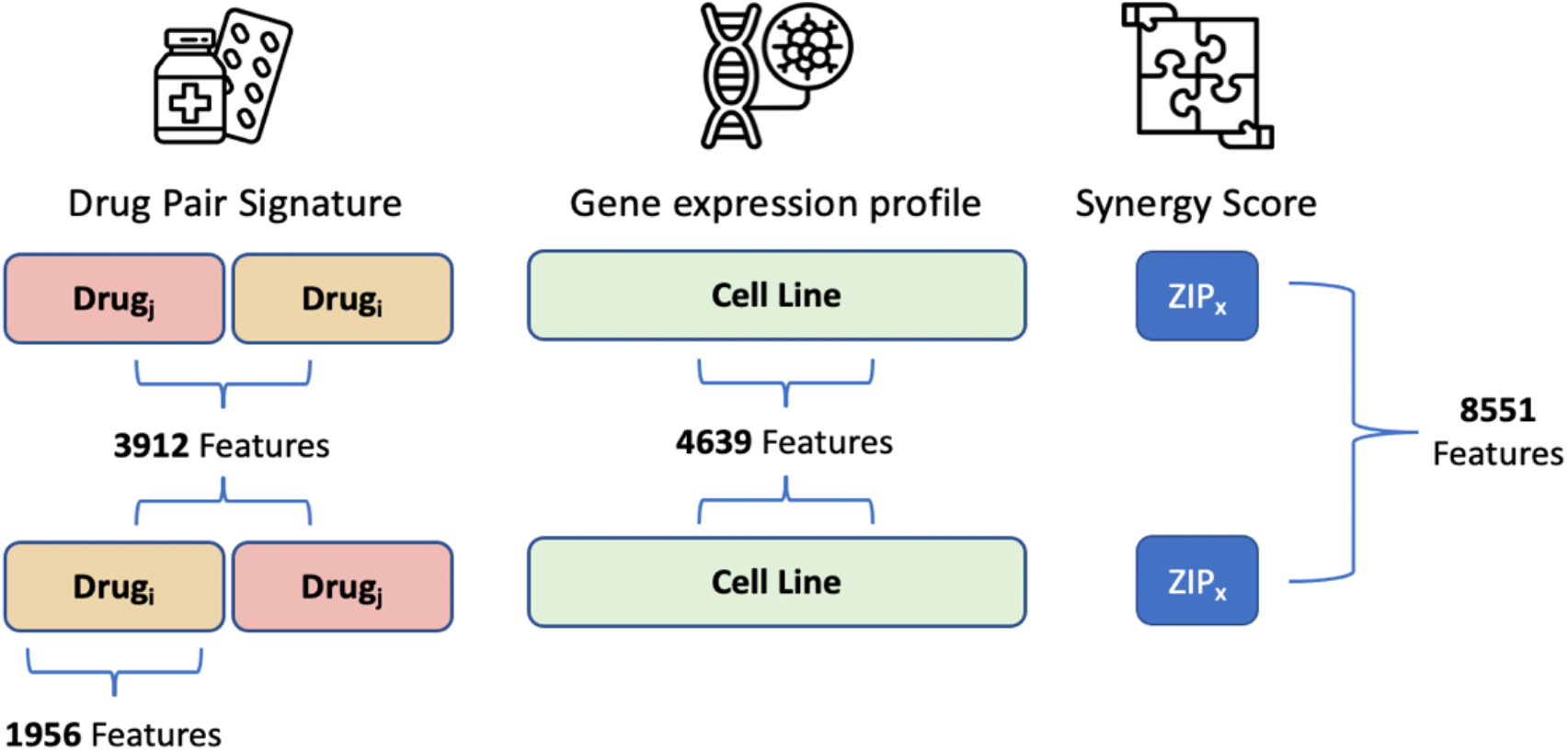
Overview of the number of features representing each element of the input. To ensure the embeddings learned by MARSY are invariant to order of the drug-pairs, for each such pair, an additional sample is included by inverting their order while keeping the same associated synergy score.

To represent drugs (as inputs to the model) we used their signatures corresponding to the differential gene expression profile of two CCLs (MCF7 and PC3) measured 24 hours after treatment with that drug with the dosage of 10 *μM*. We downloaded level 5 signatures from the Library of Integrated Network-Based Cellular Signatures (LINCS) (Subramanian, et al., 2017), and only used experimentally measured values corresponding to 978 landmark genes. The two CCLs, the time after treatment, and the drug dosage were selected to maximize the number of samples in our study. In instances where replicate signatures existed for the same CCL, same drug, same dosage, and same time-point, we calculated their mean to obtain a consensus signature. We concatenated the MCF7 and PC3 signatures of each drug, resulting in a vector of length 3912 to represent each drug-pair (Figure 2).

DrugComb is one of the largest publicly available datasets for drug synergy scores with a total of 739,964 drug combinations obtained from 8397 unique drugs. We downloaded drug synergy scores (and single drug responses) corresponding to triples of (drug1, drug2, CCL) for which we had both drug signatures (from LINCS) and CCL gene expression profiles (from CCLE) from DrugComb (V1.5, https://drugcomb.fimm.fi). Various measures of drug synergy have been introduced in the literature, which typically rely on the deviation of the observed response compared to that of a theoretical model. In this work, we focused on two more recent measures of synergism, Zero Interaction Potency (“ZIP”) and “S_mean_”, which have been introduced to overcome some of the shortcomings of older measures and provide more robust synergy scores. ZIP calculates the synergism of a drug-pair under the assumption that there is no potency between the single drug responses (Yadav, et al., 2015). ZIP incorporates both the Loewe additivity (Loewe, 1953) and the Bliss independence (Bliss, 1939) theoretical models in order to capture different types of drug interaction patterns and has been shown to be more robust and less sensitive to experimental noise (Yadav, et al., 2015). On the other hand, S_mean_ calculates synergism of a drug-pair assuming that the single drug effect is equivalent to the average of the single drug responses (Malyutina, et al., 2019). As opposed to ZIP, S_mean_ is not calculated for each combination of dosages; instead, a drug sensitivity measure is computed directly from the dose response curve of each drug and used as the drug-pair’s response. Thus, S_mean_ serves not only as a measure of drug-pair synergy but also of its sensitivity, which aims to illustrate the actual response of a CCL (Malyutina, et al., 2019), and hence is more biologically informative. For these reasons, we focused on these two measures, since they represent recent advances in defining drug synergy with several advantages compared to older models.

Since DrugComb contains (sometimes inconsistent) replicates, it is important to carefully process such cases. To identify samples with inconsistent replicates, we used two conditions. First, we only kept samples for which at least 60% of the replicates had the same synergy score sign. Second, we only kept the samples for which the standard deviation of the synergy score over replicates did not exceed 0.1. Finally, to obtain a single consensus synergy score for each sample, we calculated the replicates’ median score. The final dataset, comprised of samples that were present in all three datasets above and passed the filtering steps, contained 43,181 (drug1, drug2, CCL) triple samples corresponding to 670 unique drugs, 2353 unique drug-pairs and 75 unique CCLs. Finally, in order to ensure that the trained model is invariant to the order of the drugs in its input space, for each triple sample we constructed a new sample by changing the order of the drugs (Figure 2), resulting in a total of 86,348 triple samples. It is important to note that the samples with known synergy scores above only correspond to 0.128% of all possible combinations of 670 drugs and 75 CCLs, further emphasizing the importance of transductive computational models to complete this sparse 3-dimensional tensor.

We used relative inhibition (RI), available in DrugComb, as the response of each drug in a CCL. RI represents the normalized area under the dose-response curve after a *log*_10_ transformation (Zheng, et al., 2021). More precisely, RI is the proportion of this area under the dose-response curve to the maximal area that one drug can achieve at the same range of dosages. This metric represents the overall inhibition effect of a drug and has shown to be a robust way to represent drugs’ sensitivity (Douglass, et al., 2022). Additionally, the RI metric allows the comparison of drug responses obtained using different dose ranges (Zheng, et al., 2021).

### Baseline models

We considered several baseline models for benchmarking the performance of MARSY. Off-the-shelf models included LASSO, ElasticNet, Support Vector Machine Regression (SVM), Random Forests (RF) and a multi-layer perceptron (MLP) with the same depth and width as MARSY. State-of-the-art algorithms, developed by others for the same task, included TreeCombo (Janizek, et al., 2018), DeepSynergy (Preuer, et al., 2017), and MatchMaker (Kuru, et al., 2022). All models were trained on the same dataset (with identical folds across models) and evaluated similarly to ensure a fair comparison. Hyperparameters of these models were tuned using the same independent validation set used to tune MARSY’s hyperparameters (see Supplementary Methods for details). The model that performed best on the validation set was used for cross-validation analysis on the main dataset. In addition to the baselines above, we performed an extensive ablation study to assess the effect of each component of MARSY’s architecture on its performance. The details of the ablation study are provided in Results.

### Cross-validation, data splitting, and evaluation metrics

The performance of MARSY (and other methods) was evaluated using a five-fold cross validation (CV) approach combined with two data splitting strategies. The main dataset described above was randomly divided into five folds (based on each data split strategy described below); four folds were used to train each model using the hyperparameters obtained from the independent validation set (not used as part of these folds) and the remaining fold was used for performance evaluation. This was repeated five times so that each fold is set as the test set once. The mean and standard deviation of performance metrics on these test sets were used for evaluation.

Two data splitting strategies were used corresponding to the transductive setup of this study: leave-triple-out and leave-pair-out. In the former strategy, random triples (drug-pair and CCL) were selected randomly to construct the five folds. In the latter strategy, the folds were randomly selected based on drug-pairs, to ensure that the pairs of drugs are not seen together in the training set. To ensure a fair comparison, we used the exact same folds for all models and evaluated them in a similar manner. We evaluated the performance of MARSY and other methods in predicting the continuous-valued drug synergy scores using Root Mean Squared Error (RMSE), Spearman’s Rank Correlation Coefficient (SCC), and Pearson Correlation Coefficient (PCC). Additionally, we converted drug synergy scores to binary classes (synergistic or antagonistic) using different thresholds (0, ±1, ±5, ±10 and ±20), and used area under the receiver operating characteristic (AUROC) to evaluate the performance. For a threshold *t*, synergy scores above +*t* were considered synergistic and values below −*t* were considered antagonistic.

## RESULTS

### Performance of MARSY in prediction of drug synergy scores in a leave-triple-out data split

As the first evaluation study, we focused on the five-fold CV with leave-triple-out setup, discussed in Methods (Table 1). MARSY outperformed all baseline models using all performance metrics. In particular, MARSY achieved a Pearson correlation coefficient of PCC = 0.886 on ZIP and PCC = 0.864 on S_mean_. The substantial difference between the performance of MARSY and the MLP with a similar depth and width, reveals the importance of learning separate representations for input pairs and triples, and using a multi-task predictor to constraint these representations using single drug priors. In this evaluation, all state-of-the-art algorithms outperformed off-the-shelf methods, yet did not achieve MARSY’s performance.

**Table 1:** The performance of MARSY and baseline methods using 5-fold leave-triple-out evaluation. The folds and input data are the same across different models for a fair comparison. Best performance values are in bold-face and underlined. The mean and standard deviations are calculated across the folds. Models are sorted based on their ZIP PCC values.

| Model | ZIP | | | $S_{\text{mean}}$ | | |
| --- | --- | --- | --- | --- | --- | --- |
|  | SCC | PCC | RMSE | SCC | PCC | RMSE |
| MARSY | <u>**0.780**</u><br>($\pm 0.010$) | <u>**0.886**</u><br>($\pm 0.005$) | <u>**5.36**</u><br>($\pm 0.19$) | <u>**0.836**</u><br>($\pm 0.002$) | <u>**0.864**</u><br>($\pm 0.003$) | <u>**8.42**</u><br>($\pm 0.09$) |
| MatchMaker | 0.742<br>($\pm 0.004$) | 0.873<br>($\pm 0.006$) | 6.11<br>($\pm 0.319$) | 0.810<br>($\pm 0.003$) | 0.840<br>($\pm 0.005$) | 9.84<br>($\pm 0.23$) |
| TreeCombo<br>(XGBoost) | 0.737<br>($\pm 0.004$) | 0.870<br>($\pm 0.003$) | 5.73<br>($\pm 0.17$) | 0.817<br>($\pm 0.002$) | 0.852<br>($\pm 0.002$) | 8.77<br>($\pm 0.06$) |
| DeepSynergy | 0.701<br>($\pm 0.003$) | 0.869<br>($\pm 0.004$) | 5.78<br>($\pm 0.18$) | 0.803<br>($\pm 0.003$) | 0.843<br>($\pm 0.002$) | 9.14<br>($\pm 0.12$) |
| MLP | 0.675<br>($\pm 0.002$) | 0.840<br>($\pm 0.004$) | 6.30<br>($\pm 0.15$) | 0.781<br>($\pm 0.011$) | 0.817<br>($\pm 0.009$) | 9.72<br>($\pm 0.24$) |
| SVM | 0.689<br>($\pm 0.004$) | 0.783<br>($\pm 0.006$) | 7.65<br>($\pm 0.19$) | 0.766<br>($\pm 0.005$) | 0.779<br>($\pm 0.004$) | 10.67<br>($\pm 0.07$) |
| Random Forests | 0.419<br>($\pm 0.003$) | 0.646<br>($\pm 0.016$) | 9.04<br>($\pm 0.21$) | 0.619<br>($\pm 0.006$) | 0.640<br>($\pm 0.005$) | 12.98<br>($\pm 0.05$) |
| LASSO | 0.342<br>($\pm 0.009$) | 0.433<br>($\pm 0.013$) | 10.41<br>($\pm 0.21$) | 0.561<br>($\pm 0.011$) | 0.525<br>($\pm 0.013$) | 14.22<br>($\pm 0.14$) |
| Elastic Net | 0.342<br>($\pm 0.009$) | 0.432<br>($\pm 0.012$) | 10.42<br>($\pm 0.21$) | 0.562<br>($\pm 0.011$) | 0.525<br>($\pm 0.013$) | 14.22<br>($\pm 0.15$) |

Next, we sought to determine the ability of MARSY (and other models) in predicting the synergistic and antagonistic drugs in each CCL (a binary classification task). Since the ZIP scores and S_mean_ are continuous values, we used five thresholds (0, ±1, ±5, ±10 and ±20) with an increasing degree of strictness to binarize these values. For a specific threshold ±*t*, a score (ZIP or S_mean_) of a triple (two drugs and a CCL) larger than +*t* was considered to be synergistic, smaller than −*t* was considered to be antagonistic, and values between these two thresholds were considered low-confidence and were discarded. Figure 3A and Table S1 show the AUROC of MARSY and other baselines models as a function of these thresholds, revealing the superior performance of MARSY in prediction of ZIP score in a classification formulation.

**Figure 3:**
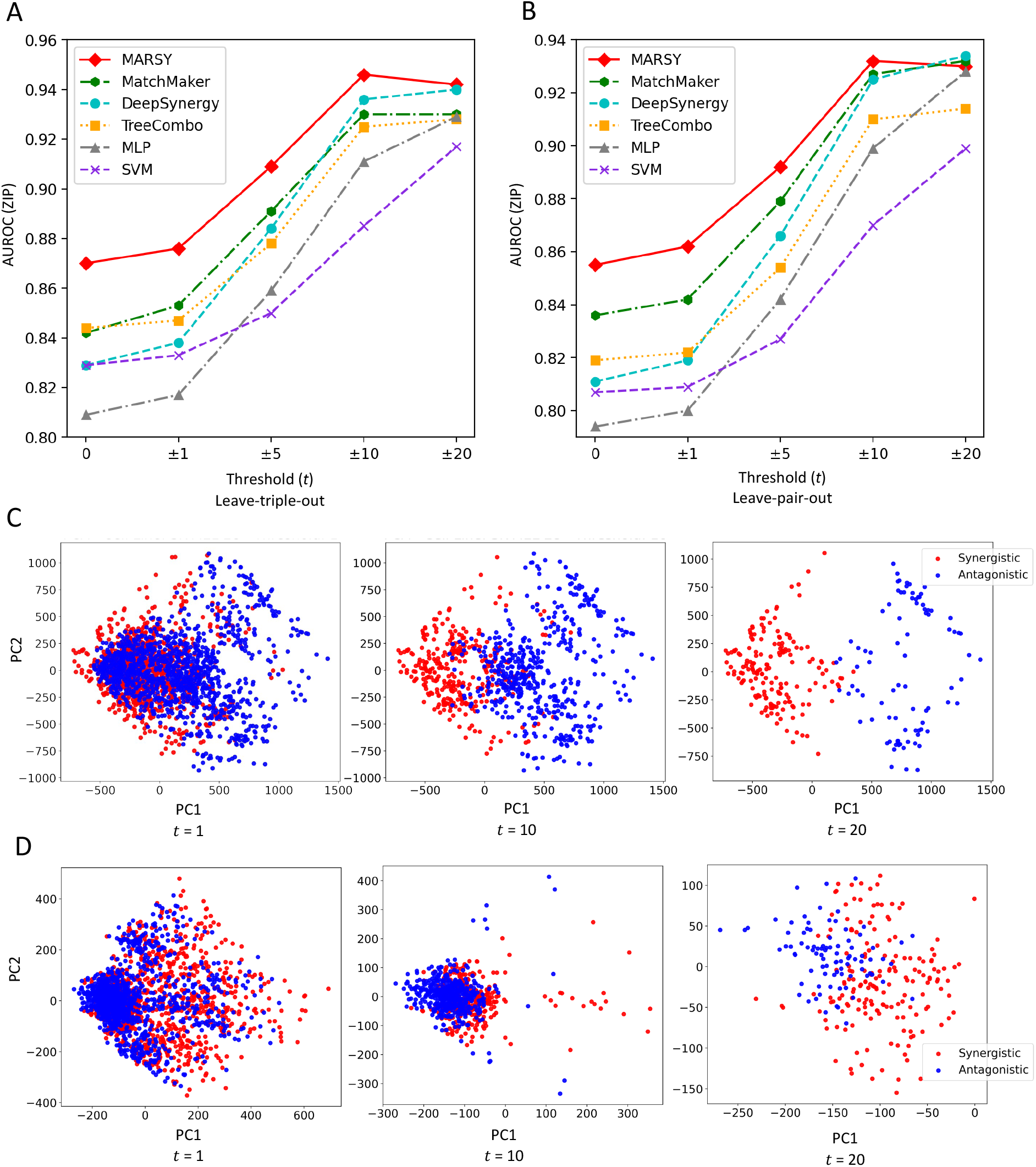
Performance of MARSY and its embeddings in identifying highly antagonistic and synergistic drugs. Panels A) and B) show classification performance of different methods in terms of AUROC for ZIP score for leave-triple-out and leave-pair-out five-fold cross validation, respectively. Panels C) and D) show principal component analysis (PCA) of synergistic and antagonistic drug-pairs for different thresholds (*t*) for SK-MEL-28 cell line based on their triple representations (panel C) and their pair representations (panel D) learned by MARSY.

### Performance of MARSY in prediction of drug synergy scores in a leave-pair-out data split

Next, we sought to evaluate the performance of MARSY, when folds in the five-fold CV are determined only based on drug-pair identities (i.e., a drug-pair in the test set is never observed in the training set). Similar to the leave-triple-out framework, MARSY outperformed all other models (Table 2), reaching a Pearson correlation coefficient value of PCC = 0.875 for prediction of ZIP score and PCC = 0.841 for prediction of S_mean_. Similar to the leave-triple-out, MatchMaker, TreeCombo and DeepSynergy were the best performing models after MARSY. Figure 3B and Table S1 show the AUROC of different models for different thresholds, further demonstrating the superior performance of MARSY for the majority of the thresholds.

**Table 2:** The performance of MARSY and baseline methods using 5-fold leave-pair-out evaluation. The folds and input data are the same across different models for a fair comparison. Best performance values are in bold-face and underlined. The mean and standard deviations are calculated across the folds. Models are sorted based on their ZIP PCC values.

| Model | ZIP | | | $S_{mean}$ | | |
| --- | --- | --- | --- | --- | --- | --- |
|  | SCC | PCC | RMSE | SCC | PCC | RMSE |
| MARSY | **<u>0.749</u>**<br>($\pm 0.011$) | **<u>0.875</u>**<br>($\pm 0.005$) | **<u>5.62</u>**<br>($\pm 0.15$) | **<u>0.809</u>**<br>($\pm 0.007$) | **<u>0.841</u>**<br>($\pm 0.011$) | **<u>9.06</u>**<br>($\pm 0.45$) |
| MatchMaker | 0.720<br>($\pm 0.006$) | 0.864<br>($\pm 0.007$) | 6.23<br>($\pm 0.12$) | 0.788<br>($\pm 0.009$) | 0.816<br>($\pm 0.015$) | 10.34<br>($\pm 0.46$) |
| TreeCombo<br>(XGBoost) | 0.689<br>($\pm 0.005$) | 0.856<br>($\pm 0.006$) | 6.00<br>($\pm 0.15$) | 0.775<br>($\pm 0.010$) | 0.815<br>($\pm 0.011$) | 9.69<br>($\pm 0.37$) |
| DeepSynergy | 0.676<br>($\pm 0.006$) | 0.860<br>($\pm 0.008$) | 5.95<br>($\pm 0.16$) | 0.762<br>($\pm 0.004$) | 0.804<br>($\pm 0.011$) | 10.17<br>($\pm 0.42$) |
| MLP | 0.642<br>($\pm 0.030$) | 0.822<br>($\pm 0.023$) | 6.68<br>($\pm 0.45$) | 0.744<br>($\pm 0.029$) | 0.752<br>($\pm 0.091$) | 11.36<br>($\pm 0.24$) |
| SVM | 0.649<br>($\pm 0.008$) | 0.773<br>($\pm 0.012$) | 7.79<br>($\pm 0.14$) | 0.737<br>($\pm 0.011$) | 0.752<br>($\pm 0.013$) | 11.15<br>($\pm 0.42$) |
| Random<br>Forests | 0.413<br>($\pm 0.005$) | 0.650<br>($\pm 0.012$) | 9.02<br>($\pm 0.17$) | 0.608<br>($\pm 0.016$) | 0.627<br>($\pm 0.013$) | 13.14<br>($\pm 0.33$) |
| LASSO | 0.333<br>($\pm 0.019$) | 0.430<br>($\pm 0.007$) | 10.43<br>($\pm 0.17$) | 0.555<br>($\pm 0.013$) | 0.519<br>($\pm 0.012$) | 14.29<br>($\pm 0.39$) |
| Elastic Net | 0.333<br>($\pm 0.018$) | 0.429<br>($\pm 0.008$) | 10.44<br>($\pm 0.18$) | 0.555<br>($\pm 0.013$) | 0.512<br>($\pm 0.019$) | 14.29<br>($\pm 0.40$) |

### Detailed evaluation of MARSY’s architecture

One aspect of MARSY’s architecture is its multitask predictor. We replaced this predictor with a single-task predictor that only predicts drug synergy scores. This modification resulted in a lower performance based on all three measures and both data split strategies (Supplementary Table S2). For example, MARSY had a 6.12% higher SCC in leave-triple-out and a 5.05% higher SCC in leave-pair-out 5-fold CV for prediction of ZIP score. These results further demonstrate the importance of including a multi-task predictor in the architecture of MARSY. In fact, changing the predictor of DL baseline models also improved their performance (Supplementary Table S2).

Next, we sought to determine the role of input representations and encoders’ architecture (Figure 1A) on MARSY’s performance. Figure 3C and 3D (visually) show that both triple and pair representations are able to separate the synergistic and antagonistic drug-pairs. To systematically evaluate their role, we implemented eight alternative architectures in which the predictor was identical to MARSY’s multitask predictor, but the latent embeddings were learned using different number and types of encoders. Supplementary Figures S1-S6 show the architecture of these models and Table 3 shows their performance. These results show that overall, MARSY’s choice of encoders perform better compared to these alternatives. In particular, using both Triple and Pair encoders is better than only using the Triple encoder (Model1 (v1)) or replacing the Triple or Pair encoders with a CCL encoder (Models 2 and 3). Similarly, it performs better compared to using separate encoders for each drug (Models 4-6).

**Table 3:** The performance of different combination of embeddings using 5-fold leave-pair-out CV. Folds are the same across different models for a fair comparison. Best performance values are in bold-face and underlined. The mean and standard deviations are calculated across the folds. Architecture of the models are provided in Figure 1 and Supplementary Figures S1-S6. In this table, Num. Enc. shows number of encoders and Num. Param shows number of parameters of each model. In this table, Model 1 (v1) uses a triple encoder identical to that of MARSY, but the encoder of Model1 (v2) and Model1 (v3) are modified such that the number of parameters of the model become close to that of MARSY.

| Model | Num. Enc. | Num. Param | Encoder Type | ZIP |  |  | S<sub>mean</sub> |  |  |
| --- | --- | --- | --- | --- | --- | --- | --- | --- | --- |
|  |  |  |  | SCC | PCC | RMSE | SCC | PCC | RMSE |
| MARSY | 2 | 61M | Pair, Triple | <u>**0.741**</u><br>(±0.018) | <u>**0.871**</u><br>(±0.009) | 5.70<br>(±0.14) | <u>**0.808**</u><br>(±0.010) | <u>**0.840**</u><br>(±0.008) | <u>**9.07**</u><br>(±0.35) |
| Model1 (v1) | 1 | 36M | Triple | 0.679<br>(±0.024) | 0.848<br>(±0.011) | 6.22<br>(±0.18) | 0.762<br>(±0.007) | 0.801<br>(±0.012) | 10.17<br>(±0.44) |
| Model1 (v2) | 1 | 62M | Triple | 0.669<br>(±0.012) | 0.838<br>(±0.006) | 6.15<br>(±0.11) | 0.768<br>(±0.006) | 0.807<br>(±0.008) | 9.97<br>(±0.38) |
| Model1 (v3) | 1 | 57M | Triple | 0.706<br>(±0.012) | 0.862<br>(±0.006) | 5.89<br>(±0.19) | 0.790<br>(±0.005) | 0.830<br>(±0.011) | 9.36<br>(±0.47) |
| Model2 | 2 | 23M | CCL, Pair | 0.706<br>(±0.018) | 0.857<br>(±0.009) | 6.52<br>(±0.13) | 0.793<br>(±0.010) | 0.826<br>(±0.007) | 10.85<br>(±0.34) |
| Model3 | 2 | 62M | CCL,<br>Triple | 0.734<br>($\pm 0.017$) | 0.867<br>($\pm 0.012$) | 5.77<br>($\pm 0.18$) | 0.804<br>($\pm 0.011$) | 0.837<br>($\pm 0.007$) | 9.17<br>($\pm 0.35$) |
| Model4 | 3 | 19M | Drug1,<br>Drug2,<br>CCL | 0.714<br>($\pm 0.019$) | 0.861<br>($\pm 0.008$) | 6.26<br>($\pm 0.15$) | 0.793<br>($\pm 0.009$) | 0.828<br>($\pm 0.007$) | 10.72<br>($\pm 0.34$) |
| Model5 | 3 | 42M | Drug1,<br>Drug2,<br>Triple | 0.737<br>($\pm 0.013$) | **0.871**<br>($\pm 0.008$) | 5.80<br>($\pm 0.15$) | 0.798<br>($\pm 0.008$) | 0.830<br>($\pm 0.009$) | 9.94<br>($\pm 0.44$) |
| Model6 | 4 | 33M | Drug1,<br>Drug2,<br>CCL,<br>Pair | 0.736<br>($\pm 0.016$) | 0.870<br>($\pm 0.008$) | **5.69**<br>($\pm 0.14$) | 0.800<br>($\pm 0.010$) | 0.833<br>($\pm 0.005$) | 9.38<br>($\pm 0.31$) |

In addition to the Model1 (v1), which uses a triple encoder with similar architecture to that of MARSY, we implemented two other variations of Model 1 with triple encoders (one with two hidden layers, but larger width and one with four hidden layers) to bring the number of their learnable parameters closer to that of MARSY (Table 3, Supplementary Figure S1). Table 3 shows that MARSY outperformed Model1 (v2), Model1 (v3), and Model3, all of which have similar number of parameters to MARSY. Moreover, MARSY outperformed DeepSynergy and MatchMaker (Table 1 and 2) that had 95 M and 59 M learnable parameters, respectively. These results suggest that the performance of MARSY is not simply an artifact of its number of parameters.

### Effect of hyperparameters and input features on MARSY’s performance

As described in Methods, several aspects of MARSY’s architecture were design choices selected a priori without hyperparameter tuning. To assess how these choices influence MARSY’s performance, we conducted a study based on leave-pair-out 5-fold CV in which we ran MARSY with 576 different combinations of multiple hyperparameters to predict the ZIP score. These hyperparameters and their options included the learning rate (0.01, 0.001, 0.0001), the optimizer (Adam, Adamax), the input activation function (Linear, ReLU), the batch size (64, 128), the dropout (with or without) and the width and depth of the encoders and the predictor (see Supplementary Table S3 in Supplementary File S1 for the specific choices).

Our analysis revealed that a large learning rate (0.01) results in poor performance with Adamax (Figure 4A) and also results in non-convergence of the Adam optimizer. However, the two smaller learning rates do not suffer from the same issue. When the learning rate is selected appropriately (the default 0.001 or 0.0001), the effects of other hyperparameters are relatively small (Figure 4B and 4C). In particular, other than the learning rate, number of encoder layers and the choice of optimizer had the largest effect, where 2 encoder hidden layers resulted in 1.4% higher median SCC (0.3% higher median PCC) compared to 3 hidden layers and Adamax resulted in 1.1% higher median SCC (0.5% higher median PCC) compared to Adam (Figure 4C and Supplementary Figure S7). In conclusion, learning rate seems to have the most effect (consistent with our experience in other related studies (Hostallero, et al., 2022)), but when large learning rates are excluded, the results are not too sensitive to the choice of hyperparameters. However, if computational complexity is not an issue, marginal improvements can be achieved using hyperparameter tuning based on an independent validation set.

**Figure 4:**
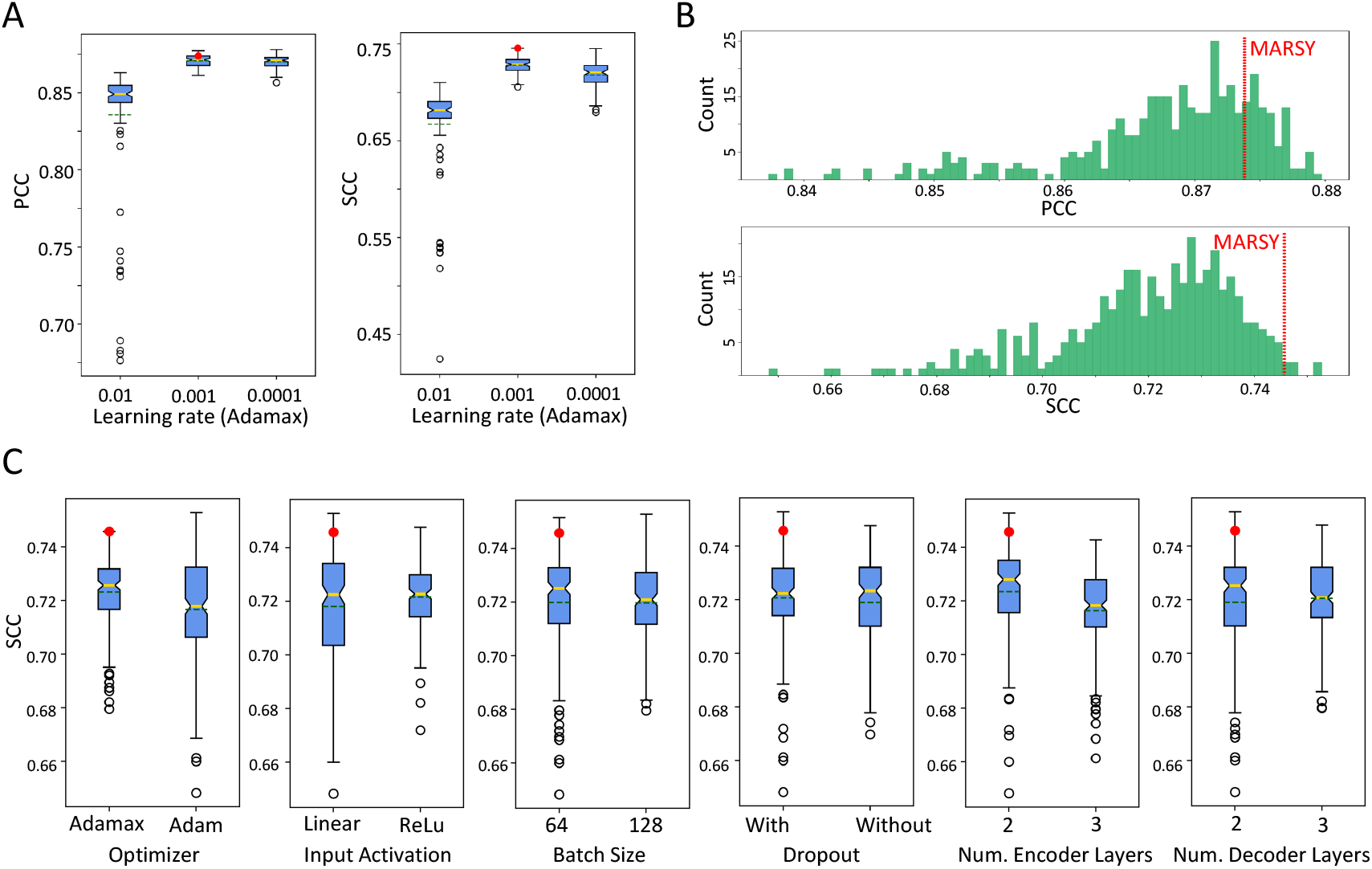
The effect of hyperparameters on performance of MARSY for prediction of ZIP score in leave-pair-out 5-fold CV. In boxplots, red circle represents MARSY, solid horizontal yellow line shows the median and dashed green line shows the mean. A) Effect of learning rate on the performance of MARSY with Adamax optimizer. B) The histogram of performance metrics based on different hyperparameter options, excluding a learning rate of 0.01. C) The boxplots show the distribution of SCC for different hyperparameter options. Runs with a learning rate of 0.01 are excluded due to the large number of non-converging runs.

MARSY uses a concatenation of LINCS drug signatures corresponding to MCF7 and PC3 cell lines. Since requiring that two LINCS drug signatures be available for each drug may be a limiting factor in applicability of MARSY, we asked whether a comparable performance can be achieved when only one of these signatures are used. Table 4 shows the performance of MARSY using 5-fold leave-pair-out cross-validation with these different options of drug signatures.

**Table 4:** The performance of MARSY using different LINCS signatures using 5-fold leave-pair-out evaluation. The folds are the same across different models for a fair comparison and are the same as the results reported in Table 2. Best performance values are in bold-face and underlined. The mean and standard deviations are calculated across the folds.

| Model | ZIP |  |  | S<sub>mean</sub> |  |  |
| --- | --- | --- | --- | --- | --- | --- |
|  | SCC | PCC | RMSE | SCC | PCC | RMSE |
| MARSY (MCF7+PC3) | <u>**0.749**</u><br>(±0.011) | <u>**0.875**</u><br>(±0.005) | <u>**5.62**</u><br>(±0.15) | <u>**0.809**</u><br>(±0.007) | <u>**0.841**</u><br>(±0.011) | <u>**9.06**</u><br>(±0.45) |
| MARSY (MCF7) | 0.742<br>(±0.012) | 0.869<br>(±0.003) | 5.75<br>(±0.14) | 0.808<br>(±0.007) | 0.840<br>(±0.011) | 9.11<br>(±0.41) |
| MARSY (PC3) | 0.742<br>(±0.012) | 0.872<br>(±0.004) | 5.68<br>(±0.16) | 0.805<br>(±0.008) | 0.836<br>(±0.013) | 9.25<br>(±0.48) |

These results show a comparable performance between using a single LINCS signature and combining the two in MARSY. For example, the SCC for MARSY with MCF7 is only 0.1% lower than the SCC for MARSY with both PC3 and MCF7 signatures in predicting S_mean_. Comparing these results with Table 2 also shows that MARSY with any of these signatures above outperforms all baseline models.

We next asked how the performance of MARSY changes if alternative drug signatures (e.g., based on their chemical structure) is used instead of LINCS signatures. We obtained the chemical representations of drugs from the DrugBank database (Wishart, et al., 2018) and used the RDKit python library (Landrum, 2006) to extract two types of chemical feature representations: “Morgan fingerprints” and “molecular descriptors”. To make the evaluation consistent, we only conserved samples from DrugComb for which we could obtain both chemical feature representations and LINCS drug signatures, along with the gene expression of the CCLs, resulting in 73,768 samples. Table 5 shows the performance of MARSY with different drug features using this dataset. Considering all three metrics on the two evaluation setups, LINCS signature performed better compared to Morgan fingerprint and molecular descriptor. We also concatenated different features to provide a combination of them as input to the model. Combination of Morgan fingerprints and molecular descriptors performed better than each of them individually in leave-triple-out, but its performance was inferior to Morgan fingerprints in leave-pair-out. Additionally, combining all three types of representations performed slightly better than LINCS signature alone (and the other representation choices), however one shortcoming of this combination is that it requires availability of multiple type of representations for each drug, which sometimes limits the number of datapoints for analysis.

The LINCS signatures used by MARSY were selected to maximize the number of training examples, which is crucial for training any DL model. However, chemical structure data (e.g., Morgan fingerprints) is readily available for a larger number of drugs, which may allow to further increase the size of the training set. On the other hand, LINCS drug signatures corresponding to other CCLs are also available, which can be used as alternatives to MCF7 and PC3. We sought to assess the trade-off between the choice of drug representations and the availability of data to form larger training sets. For this purpose and using the same data cleaning and pre-processing procedure used for forming the main dataset, we identified 6,762 triples for which Morgan fingerprints as well as LINCS molecular signatures of MCF7 (breast cancer), PC3 (prostate cancer), A549 (lung cancer), and A375 (skin cancer) were available. We used data corresponding to these triples to form our validation set (10%) and test set (90%). Then, we formed separate training sets for each type of drug representation to include all triples with data on that drug representation. This allows us to form different training set sizes depending on the availability of each type of drug representation and assess the effect of the training set size on the performance.

Performance of MARSY with different drug representations and their corresponding training set sizes are provided in Supplementary Table S4. In this analysis, Morgan fingerprints resulted in the largest training set (n = 154,718, nearly double the size of the training set of MCF7 that had the second largest training set), but its performance was not the best in any of the categories. However, in most categories, it was only less than 2% worse than the best performing option. A375 signatures resulted in the worst performance, but this was expected due to its extremely small training set size of only 1,558 samples. Although A549’s training set was ^~^32% smaller than that of MCF7, it still resulted in acceptable predictions (e.g., S_mean_ PCC = 0.852). These results suggest that while extremely small training set sizes affect the performance, all three LINCS signatures that had more than 50,000 training samples resulted in acceptable performance metrics. Additionally, in applications where LINCS signatures are not available, Morgan fingerprints can be used as alternatives.

Next, we asked how the performance of MARSY compares against DeepSynergy, MatchMaker, and TreeCombo, when Morgan fingerprints are used as drug representations. Supplementary Table S5 shows the performance of these models using the same training, validation, and test sets described above. These results show that even when Morgan fingerprints are used, MARSY outperforms these baseline models in most categories.

In a recent study, a high degree of correlation between the synergy of drug pairs and the correlation of the transcriptomic profiles of the corresponding monotherapies in the same cell line was observed, which motivated a drug synergy score predictor based on this principle (Diaz, et al., 2020). We designed a small study to investigate whether we also observe a correlation between the monotherapy LINCS transcriptomic signatures and their synergism in the same CCLs. We focused on three cell lines MCF7, PC3, and A549 for which we found a large number of signatures and we used in our LINCS signature analysis discussed above. For each (CCL, drug1, drug2) triple in our dataset (where the CCL is one of the three CCLs above), we calculated the correlation between the LINCS signature of drug1 and drug2 in that CCL. We then calculated the correlation of these values with the synergy score of those drugs in that CCL. Based on these analyses, we did not observe a similar pattern to that reported in (Diaz, et al., 2020) and the highest correlation values (achieved in A549 for S_mean_), were only equal to PCC = 0.08 and SCC = 0.104.

### Prediction of drug synergy scores for new triples not present in DrugComb

Next, we used our trained model to predict the ZIP synergy scores of 133,722 new (drug-pair, CCL) triples that were not present in DrugComb. These triples correspond to all possible combinations of 69 unique drugs that appeared in at least 10% of the drug-pairs in our training set (Supplementary Table S6). Given these synergy scores, we first asked which drug-pairs show a synergistic effect on all 75 CCLs. We identified seven drug-pairs that had a minimum ZIP score larger than 2 across all CCLs. The majority of these corresponded to the combination of vincristine (a chemotherapy agent) with other drugs. In particular, two of these combinations corresponded to tyrosine kinase inhibitors (TKI) lapatinib (mean ZIP = 28.6) and imatinib (mean ZIP = 17.9) combined with vincristine. Several independent studies have shown the synergistic effect of these drugs. For example, these two TKIs sensitized KBV20C oral cancer cells to vincristine, and their combination up-regulated apoptosis and reduced cell viability (KIM, et al., 2019). In addition, lapatinib significantly increased the efficacy of vincristine in epidermoid carcinoma C-A120 cells that overexpress multidrug resistance–associated protein 1 (MRP1) (Ma, et al., 2014). Another noteworthy combination corresponded to docetaxel (chemotherapy) and veliparib (a PARP inhibitor), (mean ZIP = 16.3). The siRNA knockdown of PARP1 in PC3 cell lines has been shown to enhance docetaxel activity (Wu, et al., 2013), and MARSY predicted the ZIP score of docetaxel and PARP-inhibitor veliparib in PC3 to be 9.8 (reflecting synergism).

Next, we sought to identify drug combinations that show synergistic effect in a tissue-specific manner. For this purpose, we focused on breast tissue, since it had the highest number of CCLs in our cohort (n = 26). To identify breast cancer-specific synergistic combinations, we required a drug-pair to not be antagonistic in any of the CCLs of breast cancer (minimum ZIP in the tissue > 0) and then ranked the remaining drug-pairs based on the difference between their mean ZIP score in CCLs of breast cancer and the mean ZIP score of CCLs of other tissues (Supplementary Table S7). The top-ranked drug-pair corresponded to combination of paclitaxel (a chemotherapy widely used in treating breast cancer) and ruxolitinib (a selective JAK1/2 kinase inhibitor). A one-sided Mann Whitney U test also showed this difference to be statistically significant (Benjamini-Hochberg false discovery rate = 7.76E-3). Ruxolitinib has been shown to enhance the efficacy of paclitaxel in a synergistic manner in ovarian cancer cells (Han, et al., 2018). Recent studies of triple negative breast cancer (TNBC) samples has shown that inhibition of JAK1/2 by ruxolitinib sensitizes cancer cells to paclitaxel, both *in-vitro* and *in-vivo* (Han, et al., 2021; Lian, et al., 2020). A phase I clinical trial of the combination of these two drugs in HER2-negative metastatic breast cancer patients was recently completed and showed these two drugs to be well tolerated by the participants (Lynce, et al., 2021). Following these results, a phase II randomized clinical trial is currently undergoing for the combination of these two drugs in TNBC patients (Lynce, et al., 2021). Our results also showed that TNBC CCLs in our cohort have a higher mean ZIP score for this drug combination compared to other subtypes of breast cancer (difference mean ZIP = 1.4).

These examples illustrate the utility of MARSY to suggest synergistic drug combination candidates for follow-up drug screening experiments.

## Discussion and Conclusion

In this study, we proposed MARSY, a novel deep multi-task learning method for prediction of synergy scores of drug-pairs in difference cancer cell lines. Extensive evaluations using four different metrics and two methods of data splitting for cross-validation revealed the better performance of MARSY compared to various computational models designed for this task. MARSY’s architecture was designed to learn distinct embeddings that capture different and complimentary views of the input features and are informative not just for drug synergy score prediction, but also enable single drug response prediction. Given the gene expression profile of CCLs as well as the LINCS signatures of each drug, the two encoders of MARSY learn different views of the input features, capturing the interplay of the drugs, as well as the drugs and the CCL. Our ablation study confirmed that incorporating both of these encoders is beneficial to the performance of the model. Moreover, the predictor of MARSY utilizes auxiliary tasks of single drug prediction to enhance the synergy score prediction performance, by enhancing the embeddings learned during the end-to-end training of the model. We also observed that these auxiliary tasks can be helpful to other DL-based baseline models.

Unlike many previous models that utilize drug molecular profiles, we used the signatures of each drug representing the changes in the expression profiles of two cancer cell lines after administration of the drug. We observed that using only one of these signatures results in comparable performance to that of using both signatures simultaneously (e.g., only a 0.1% difference in Spearman’s correlation in predicting S_mean_ when using only MCF7 signature, Table 4). This suggests that MARSY can also be used with only one drug signature, as long as there are a large number of samples available for training. In fact, we observed that LINCS drug signatures corresponding to different cancer types (MCF7, PC3, A549) result in comparable performance (Table S4), even though the number of training examples when using A549 drug signatures was 32% smaller than MCF7. However, when the number of training examples was very small in the case of A375, a significant deterioration of performance was observed. Our analyses using a fixed test set, but different training sets that included all samples with available drug features of each type, showed that although Morgan fingerprints were not the best performing option for MARSY (in spite of having a training set with almost double the size of largest LINCS signature), they resulted in prediction performances that were only less than 2% lower than the best performing option in most cases. Since Morgan fingerprints are available for a larger number of drugs and the use of them with MARSY results in only a small performance deterioration (Table 5 and S5), they are viable alternatives to LINCS signatures when such signatures are unavailable.

**Table 5:** The performance of MARSY using distinct types of drug features. Evaluation is performed using 5-fold leave-triple-out and 5-fold leave-pair-out on the prediction of the ZIP synergy score. Best performance values are in bold-face and underlined. The mean and standard deviations are calculated across the folds.

| Drug Features | Leave-Triple-Out |  |  | Leave-Pair-Out |  |  |
| --- | --- | --- | --- | --- | --- | --- |
|  | SCC | PCC | RMSE | SCC | PCC | RMSE |
| Molecular descriptors | 0.702<br>($\pm 0.014$) | 0.849<br>($\pm 0.005$) | 5.57<br>($\pm 0.12$) | 0.676<br>($\pm 0.016$) | 0.841<br>($\pm 0.009$) | 5.67<br>($\pm 0.35$) |
| Morgan fingerprints | 0.739<br>($\pm 0.001$) | 0.858<br>($\pm 0.007$) | 5.24<br>($\pm 0.13$) | **<u>0.729</u>**<br>($\pm 0.011$) | 0.854<br>($\pm 0.009$) | 5.36<br>($\pm 0.21$) |
| Molecular descriptors &<br>Morgan fingerprints | 0.746<br>($\pm 0.003$) | 0.861<br>($\pm 0.007$) | 5.16<br>($\pm 0.15$) | 0.714<br>($\pm 0.016$) | 0.846<br>($\pm 0.012$) | 5.42<br>($\pm 0.35$) |
| LINCS signature | 0.756<br>($\pm 0.007$) | 0.869<br>($\pm 0.005$) | 5.02<br>($\pm 0.11$) | 0.725<br>($\pm 0.012$) | 0.857<br>($\pm 0.009$) | **<u>5.22</u>**<br>($\pm 0.29$) |
| All features combined | **<u>0.757</u>**<br>($\pm 0.005$) | **<u>0.870</u>**<br>($\pm 0.005$) | **<u>5.01</u>**<br>($\pm 0.13$) | 0.727<br>($\pm 0.010$) | **<u>0.858</u>**<br>($\pm 0.008$) | **<u>5.22</u>**<br>($\pm 0.28$) |

In this study, our focus was on developing a transductive model to impute the missing values in our dataset. As such, we used MARSY to predict the synergy scores of 133,722 new (drug-pair, CCL) triples that were not present in DrugComb and validated some of these predictions and insights obtained from them using independent datasets. However, one of the remaining challenges in this domain is developing models that can predict the synergy scores for unseen CCLs. Currently, the main challenge in achieving this goal is data availability. For example, after data cleaning, pre-processing and quality control, we ended up with only 75 CCLs that were usable for method development in this study. Such small numbers do not allow deep learning model development that generalizes well to unseen CCLs. However, as new data becomes available, we expect that this shortcoming of current datasets will be resolved and accurate models that can generalize to unseen CCLs can be developed.

## Funding

This work was supported by the Government of Canada’s New Frontiers in Research Fund (NFRF) [NFRFE-2019-01290] (AE), by Natural Sciences and Engineering Research Council of Canada (NSERC) grant RGPIN-2019-04460 (AE), and by McGill Initiative in Computational Medicine (MiCM) (AE). This work was also funded by Génome Québec, the Ministère de l’Economie et de l’Innovation du Québec, IVADO, the Canada First Research Excellence Fund and Oncopole, which receives funding from Merck Canada Inc. and the Fonds de Recherche du Québec – Santé (AE). This research was enabled in part by support provided by Calcul Québec (www.calculquebec.ca) and Compute Canada (www.computecanada.ca).

## Data and Code Availability

All data generated as part of this study are provided as supplementary files. An implementation of the algorithms in Python and cleaned input datasets are freely available at: https://github.com/Emad-COMBINE-lab/MARSY.

## Supplementary Files

**Supplementary File S1:** This file contains Supplementary Methods, Supplementary Figures S1-S7, Supplementary Tables S3-S5, and the caption of Supplementary Tables S1, S2, S6, and S7, which are provided as separate xlsx files.

## Notes

### Competing Interest Statement

The authors have declared no competing interest.

### Summary of Updates

1) New analyses are added to fully characterize the effect of drug representations. 2) Minor modifications and revisions are done to improve the clarity of the manuscript.

## REFERENCES

Amzallag, A., Ramaswamy, S. and Benes, C.H. Statistical assessment and visualization of synergies for large-scale sparse drug combination datasets. BMC Bioinformatics 2019;20(1):83.

Bansal, M., et al. A community computational challenge to predict the activity of pairs of compounds. Nat Biotechnol 2014;32(12):1213–1222.

Barretina, J., et al. The Cancer Cell Line Encyclopedia enables predictive modelling of anticancer drug sensitivity. Nature 2012;483(7391):603–607.

Bayat Mokhtari, R., et al. Combination therapy in combating cancer. Oncotarget 2017;8(23):38022–38043.

Bliss, C.I. The toxicity of poisons applied jointly 1. Annals of applied biology 1939;26(3):585–615.

Celebi, R., et al. In-silico Prediction of Synergistic Anti-Cancer Drug Combinations Using Multi-omics Data. Scientific Reports 2019;9(1):8949.

Costello, J.C., et al. A community effort to assess and improve drug sensitivity prediction algorithms. Nat Biotechnol 2014;32(12):1202–1212.

Diaz, J.E., et al. The transcriptomic response of cells to a drug combination is more than the sum of the responses to the monotherapies. Elife 2020;9.

Douglass, E.F., et al. A community challenge for a pancancer drug mechanism of action inference from perturbational profile data. Cell Reports Medicine 2022;3(1):100492.

Han, E.S., et al. Ruxolitinib synergistically enhances the anti-tumor activity of paclitaxel in human ovarian cancer. Oncotarget 2018;9(36).

Han, J., et al. JAK2 regulates paclitaxel resistance in triple negative breast cancers. Journal of Molecular Medicine 2021;99(12):1783–1795.

Hostallero, D.E., Li, Y. and Emad, A. Looking at the BiG picture: incorporating bipartite graphs in drug response prediction. Bioinformatics 2022;38(14):3609–3620.

Hostallero, D.E., et al. A Deep Learning Framework for Prediction of Clinical Drug Response of Cancer Patients and Identification of Drug Sensitivity Biomarkers using Preclinical Samples. bioRxiv 2021;bioRxiv 2021.07.06.451273.

Huang, E.W., et al. Tissue-guided LASSO for prediction of clinical drug response using preclinical samples. PLoS Comput Biol 2020;16(1):e1007607.

Janizek, J.D., Celik, S. and Lee, S.-I. Explainable machine learning prediction of synergistic drug combinations for precision cancer medicine. bioRxiv 2018;bioRxiv 331769.

Jeon, M., et al. In silico drug combination discovery for personalized cancer therapy. BMC Syst Biol 2018;12(Suppl 2):16.

Kim, J.Y., Kim, H.S. and Yoon, S. Tyrosine Kinase Inhibitors Imatinib and Erlotinib Increase Apoptosis of Antimitotic Drug-resistant KBV20C Cells Without Inhibiting P-gp. Anticancer Research 2019;39(7):3785–3793.

Kingma, D.P. and Ba, J. Adam: A method for stochastic optimization. arXiv preprint arXiv:1412.6980 2014.

Kuru, H.I., Tastan, O. and Cicek, A.E. MatchMaker: A Deep Learning Framework for Drug Synergy Prediction. IEEE/ACM Trans Comput Biol Bioinform 2022;19(4):2334–2344.

Landrum, G. RDKit: Open-source Cheminformatics. In, http://Www.Rdkit.Org/. 2006.

Li, H., et al. TAIJI: approaching experimental replicates-level accuracy for drug synergy prediction. Bioinformatics 2018;35(13):2338–2339.

Li, H., et al. Network Propagation Predicts Drug Synergy in Cancers. Cancer Research 2018;78(18):5446–5457.

Li, J., et al. A Machine Learning Method for Drug Combination Prediction. Frontiers in Genetics 2020;11(1000).

Li, S., et al. Prediction of Synergistic Drug Combinations for Prostate Cancer by Transcriptomic and Network Characteristics. Frontiers in Pharmacology 2021;12(315).

Lian, B., et al. Truncated HDAC9 identified by integrated genome-wide screen as the key modulator for paclitaxel resistance in triple-negative breast cancer. Theranostics 2020;10(24):11092–11109.

Loewe, S. The problem of synergism and antagonism of combined drugs. Arzneimittelforschung 1953;3(6):285–290.

Lynce, F., et al. Phase I study of JAK1/2 inhibitor ruxolitinib with weekly paclitaxel for the treatment of HER2-negative metastatic breast cancer. Cancer Chemotherapy and Pharmacology 2021;87(5):673–679.

Ma, S.-l., et al. Lapatinib antagonizes multidrug resistance-associated protein 1-mediated multidrug resistance by inhibiting its transport function. Mol Med 2014;20(1):390–399.

Madani Tonekaboni, S.A., et al. Predictive approaches for drug combination discovery in cancer. Brief Bioinform 2018;19(2):263–276.

Malyutina, A., et al. Drug combination sensitivity scoring facilitates the discovery of synergistic and efficacious drug combinations in cancer. PLOS Computational Biology 2019;15(5):e1006752.

Preuer, K., et al. DeepSynergy: predicting anti-cancer drug synergy with Deep Learning. Bioinformatics 2017;34(9):1538–1546.

Reinhold, W.C., et al. CellMiner: A Web-Based Suite of Genomic and Pharmacologic Tools to Explore Transcript and Drug Patterns in the NCI-60 Cell Line Set. Cancer Research 2012;72(14):3499–3511.

Shankavaram, U.T., et al. CellMiner: a relational database and query tool for the NCI-60 cancer cell lines. BMC Genomics 2009;10(1):277.

Sidorov, P., et al. Predicting Synergism of Cancer Drug Combinations Using NCI-ALMANAC Data. Frontiers in Chemistry 2019;7(509).

Subramanian, A., et al. A Next Generation Connectivity Map: L1000 Platform and the First 1,000,000 Profiles. Cell 2017;171(6):1437–1452.e1417.

Sun, Y., et al. Combining genomic and network characteristics for extended capability in predicting synergistic drugs for cancer. Nature Communications 2015;6(1):8481.

Wishart, D.S., et al. DrugBank 5.0: a major update to the DrugBank database for 2018. Nucleic Acids Res 2018;46(D1):D1074–d1082.

Wu, W., et al. Inhibition of PARP1 by small interfering RNA enhances docetaxel activity against human prostate cancer PC3 cells. Biochemical and Biophysical Research Communications 2013;442(1):127–132.

Yadav, B., et al. Searching for Drug Synergy in Complex Dose–Response Landscapes Using an Interaction Potency Model. Computational and Structural Biotechnology Journal 2015;13:504–513.

Yang, W., et al. Genomics of Drug Sensitivity in Cancer (GDSC): a resource for therapeutic biomarker discovery in cancer cells. Nucleic Acids Res 2013;41(Database issue):D955–961.

Zagidullin, B., et al. DrugComb: an integrative cancer drug combination data portal. Nucleic Acids Res 2019;47(W1):W43–w51.

Zheng, S., et al. DrugComb update: a more comprehensive drug sensitivity data repository and analysis portal. Nucleic Acids Research 2021;49(W1):W174–W184.

