## Supplementary material for "MARSY: A multitask deep learning framework for prediction of drug combination synergy scores": File S1

### Supplementary Methods

#### Hyperparameter tuning and training of MatchMaker, DeepSynergy and TreeCombo

To ensure a fair and consistent comparison between baseline models and MARSY, we used the same training/validation/test sets for all models. The validation set was kept independent of the dataset used for cross-validation to ensure no data leakage occurs. For each baseline model, we performed hyperparameter tuning using the validation set in a grid format. The range and options for hyperparameters were selected based on a combination of recommendations from the original studies and initial tests on our dataset to exclude options that result in non-convergence or perform significantly worse than other options. For DeepSynergy, the final hyperparameters and their options that were used in the grid search included input activation function [ReLu, Linear], learning rate [0.0001, 0.00001], number of hidden layers [2, 3] and width of hidden layers [[8192, 4096], [4096, 8192], [8192, 4096, 2048], [4096, 8192, 4096]]. For MatchMaker, the hyperparameters and their options used in the grid form included input activation function [ReLu, Linear], learning rate [0.001, 0.0001], number of encoder layers [2, 3], number of decoder layers [2, 3], and width of layers [2048, 4096, 2048, 1024], [1024, 2048, 1024, 512], [2048, 4096, 2048, 1024, 512], [1024, 2048, 1024, 512, 256], [2048, 4096, 2048, 2048, 1024], [1024, 2048, 1024, 1024, 512], [2048, 4096, 2048, 2048, 1024, 512], [1024, 2048, 1024, 1024, 512, 256]]. For Treecombo, the hyperparameters and their options used in the grid form included number of estimators [500, 1000], maximum depth [4, 8], and learning rate [0.1, 0.01].

### Supplementary Figures

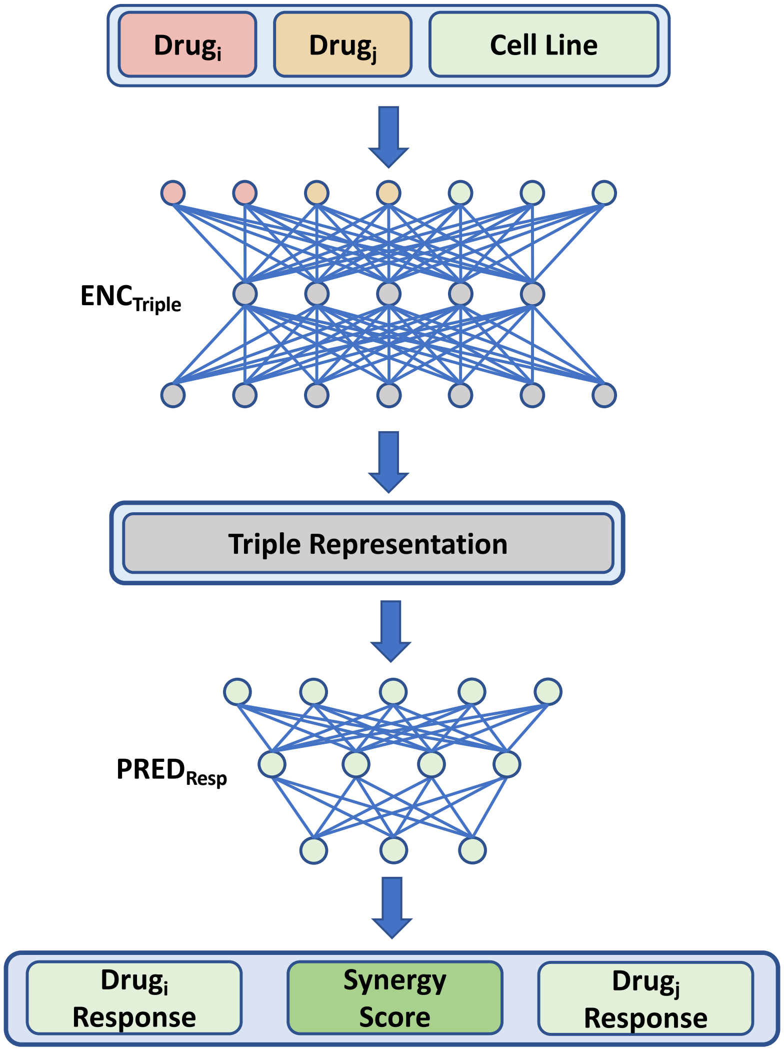


**Figure S1:** Architecture of Model1 (related to Table 3). The predictor has the same architecture as that of MARSY's. In Model1 (v1), the encoder includes two hidden layers with the following number of neurons [2048, 4096]. In Model1 (v2), the encoder includes two hidden layers with the following number of neurons [4096, 4096]. In Model1 (v3), the encoder includes four hidden layers with the following number of neurons [2048, 4096, 4096, 2048].

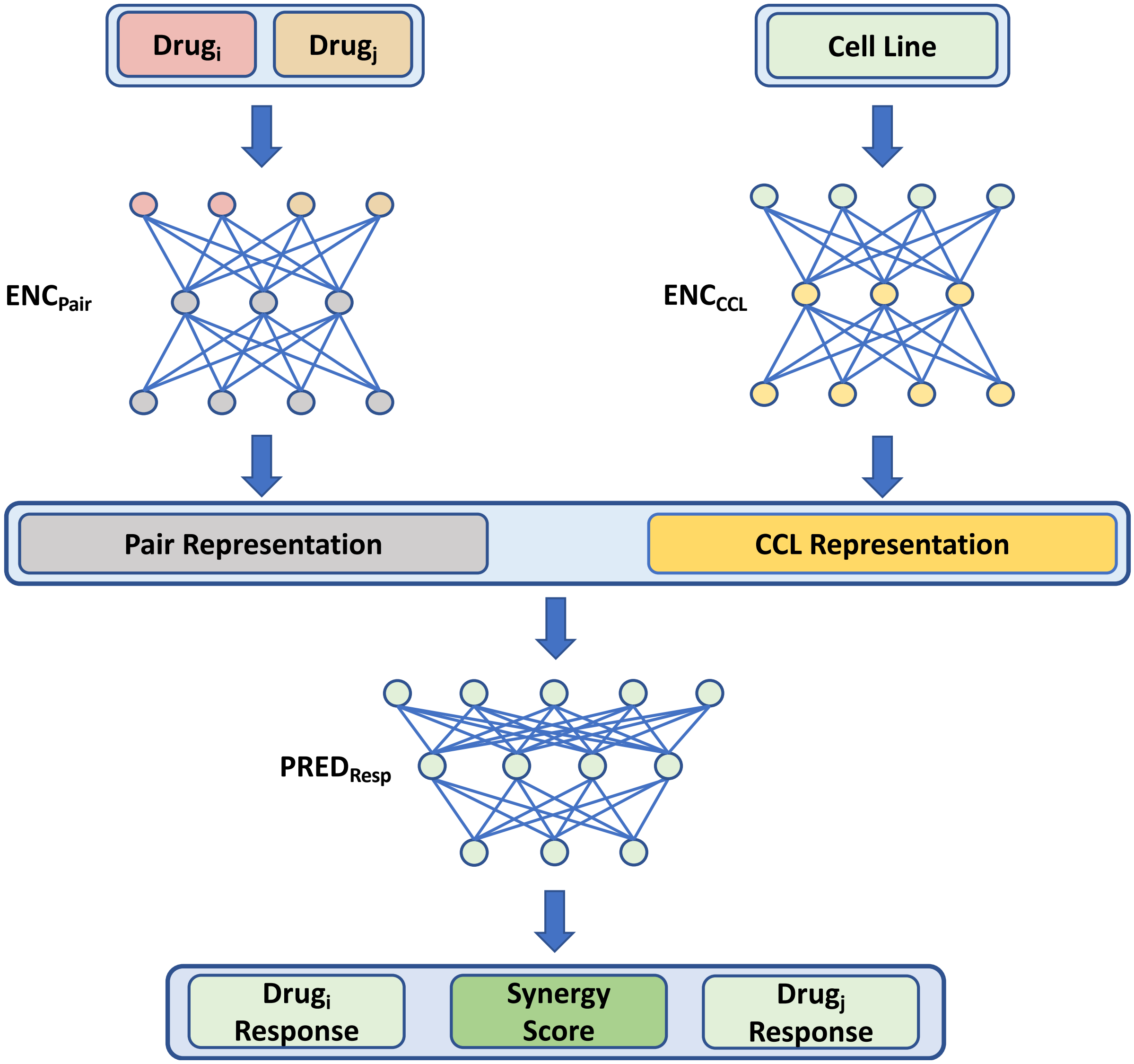


**Figure S2:** Architecture of Model2 (related to Table 3). The Pair encoder and the predictor have the same architecture as their counterparts in MARSY. CCL encoder includes 2 hidden layers with the following number of neurons [1024, 2048].

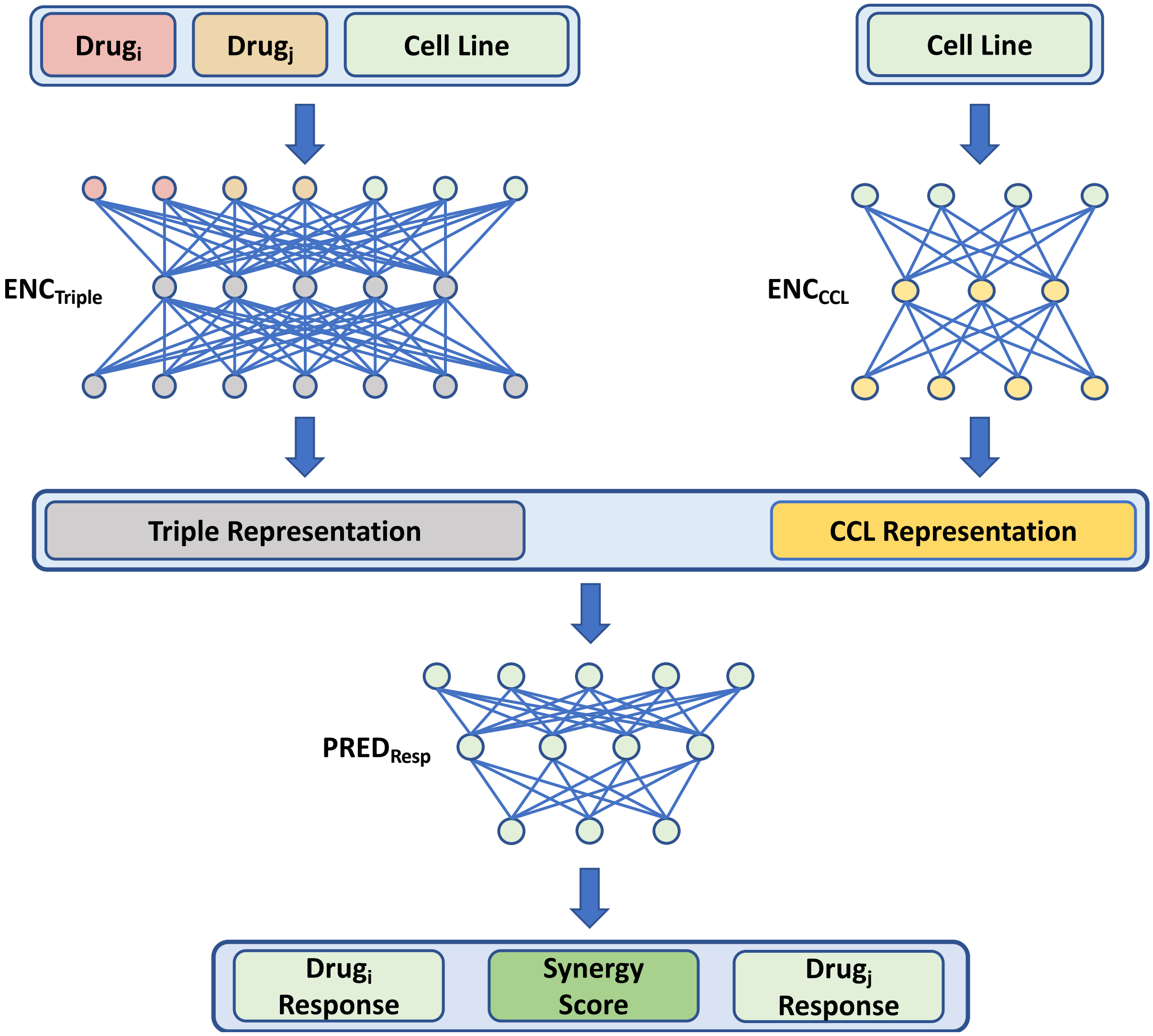


**Figure S3:** Architecture of Model3 (related to Table 3). The Triple encoder and the predictor have the same architecture as their counterparts in MARSY. CCL encoder includes 2 hidden layers with the following number of neurons [1024, 2048].

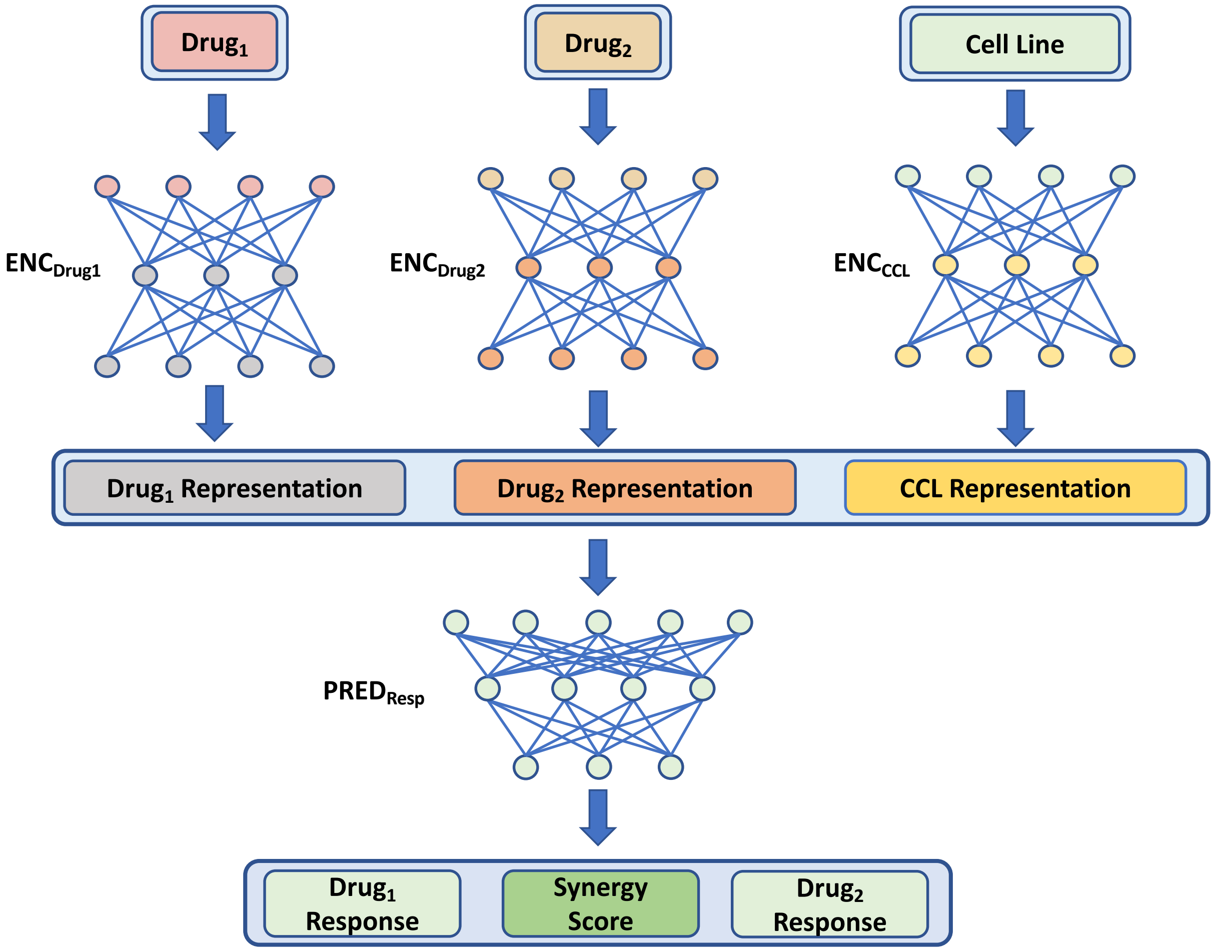


**Figure S4:** Architecture of Model4 (related to Table 3). The decoder has the same arthitecture as its counterpart in MARSY. CCL encoder includes 2 hidden layers with the following number of neurons [1024, 2048]. Each drug encoder includes 2 hidden layers with the following number of neurons [512, 1024].

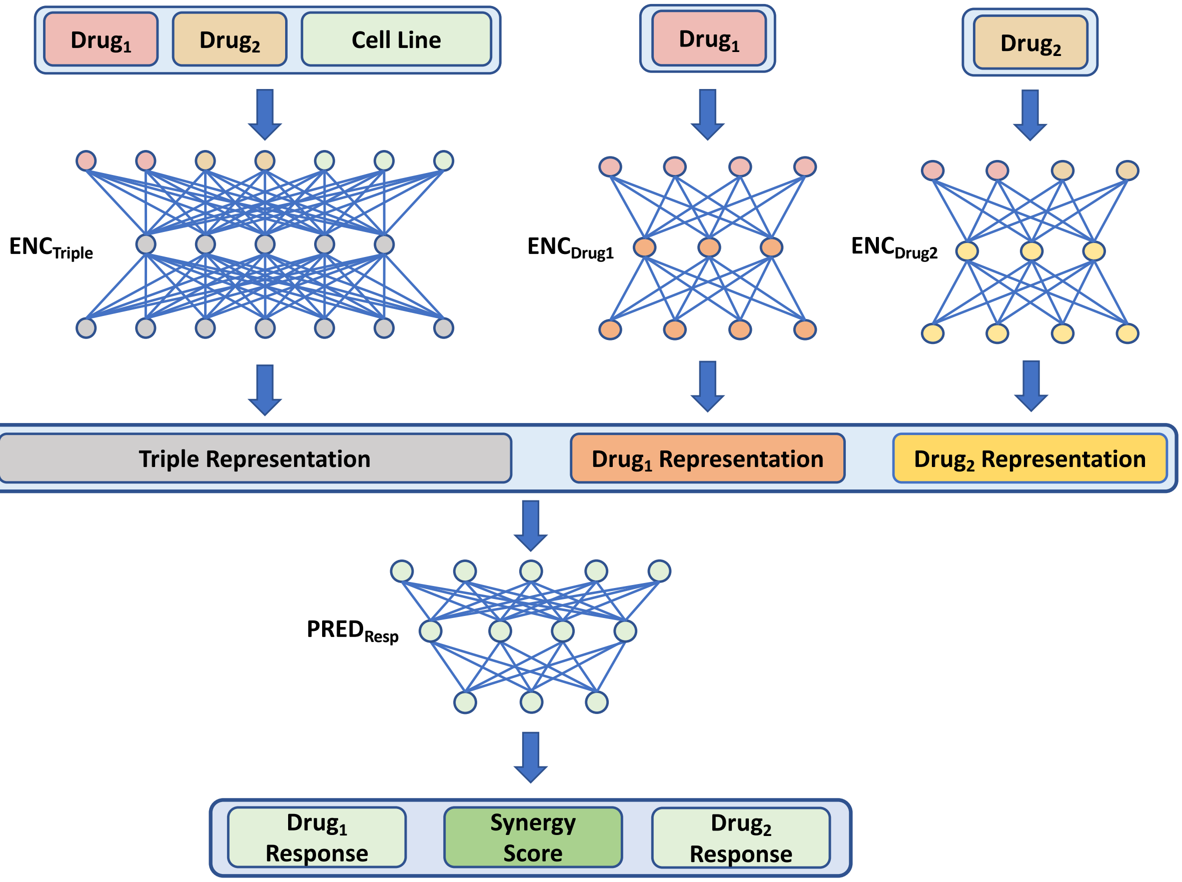


**Figure S5:** Architecture of Model5 (related to Table 3). The Triple encoder and the predictor have the same architecture as their counterparts in MARSY. Drug encoders include 2 hidden layers with the following number of neurons [512, 1024].

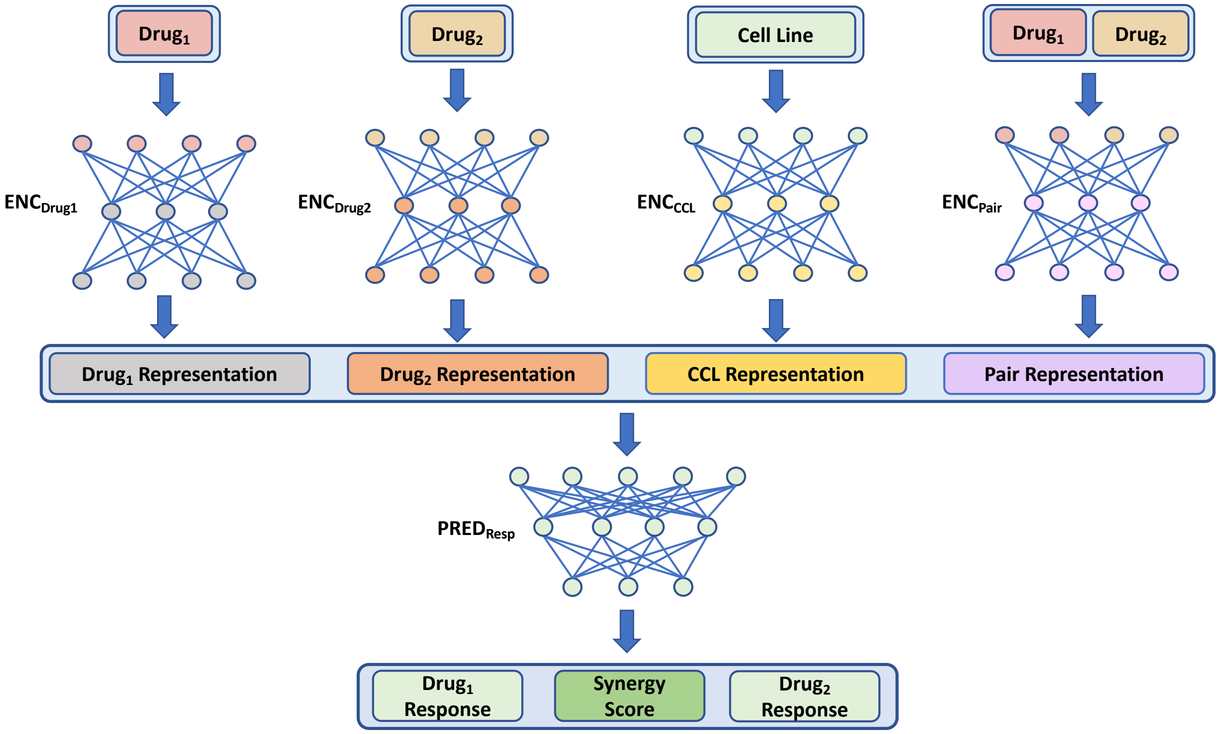


**Figure S6:** Architecture of Model6 (related to Table 3). The Pair encoder and the predictor have the same architecture as their counterparts in MARSY. CCL encoder includes 2 layers with the following number of neurons [1024, 2048]. Each drug encoder includes 2 layers with the following number of neurons [512, 1024].

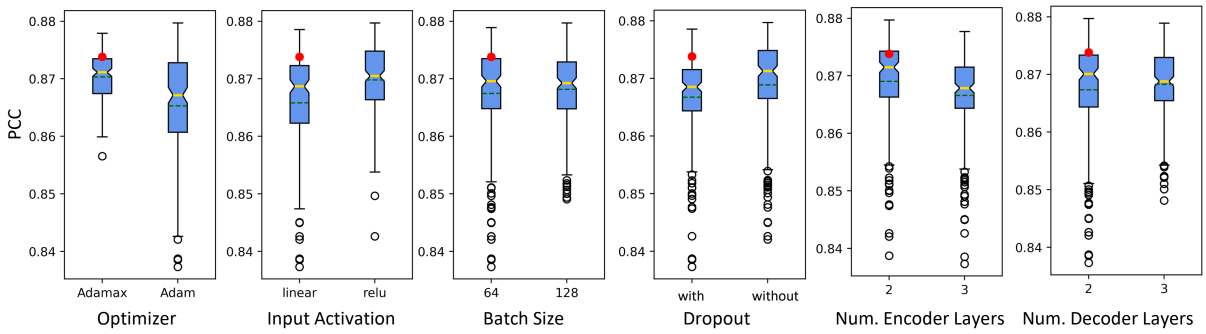


**Figure S7:** The effect of hyperparameters on performance of MARSY for prediction of ZIP score in leave-pair-out 5-fold CV. In boxplots, red circle represents MARSY, solid yellow line shows the median and dashed green line shows the mean. The boxplots show the distribution of PCC for different hyperparameter options. Runs with learning rate of 0.01 are excluded due to the large number of non-converging runs.

### Supplementary Tables

**Table S1:** AUROC of MARSY and baseline models using different thresholds. The table is provided as a separate xlsx file.

**Table S2:** The comparison between a single-task predictor and a multi-task predictor for MARSAY and other deep learning models. The table is provided as a separate xlsx file.

**Table S3:** Architectural options for the hyperparameter effects analysis. We varied the number of layers in both encoders ($ENC_{Triple}$ and $ENC_{Pair}$) and the predictor ($PRED_{Resp}$). For each option, we tested 3 different combinations of layers' width. The number of neurons chosen for each option over the different number of layers follow the same design pattern.

| Name | $ENC_{Triple}$ | | $ENC_{Pair}$ | | $PRED_{Resp}$ | |
|---|---|---|---|---|---|---|
| | Layers | Number of Neurons | Layers | Number of Neurons | Layers | Number of Neurons |
| Arch1 | 2 | [2048, 4096] | 2 | [1024, 2048] | 2 | [4096, 1024] |
| Arch2 | | [1024, 2048] | | [512, 1024] | | [2048, 512] |
| Arch3 | | [4096, 2048] | | [2048, 1024] | | [2048, 512] |
| Arch4 | 2 | [2048, 4096] | 2 | [1024, 2048] | 3 | [4096, 1024, 256] |
| Arch5 | | [1024, 2048] | | [512, 1024] | | [2048, 512, 128] |
| Arch6 | | [4096, 2048] | | [2048, 1024] | | [2048, 512, 128] |
| Arch7 | 3 | [2048, 4096, 2048] | 3 | [1024, 2048, 1024] | 2 | [2048, 512] |
| Arch8 | | [1024, 2048, 1024] | | [512, 1024, 512] | | [1024, 256] |
| Arch9 | | [4096, 2048, 1024] | | [2048, 1024, 512] | | [1024, 256] |
| Arch10 | 3 | [2048, 4096, 2048] | 3 | [1024, 2048, 1024] | 3 | [2048, 512, 128] |
| Arch11 | | [1024, 2048, 1024] | | [512, 1024, 512] | | [1024, 256, 64] |
| Arch12 | | [4096, 2048, 1024] | | [2048, 1024, 512] | | [1024, 256, 64] |

**Table S4:** The performance of MARSY using different LINCS signatures. First, we identified 6,762 triples for which Morgan fingerprints (MFP) and LINCS molecular signatures of MCF7, PC3, A549, and A375 were available. We used data corresponding to these triples to form our validation set (10%) and test set (90%). Then, we formed separate training sets for each type of drug representation to include all triples with data on that drug representation. The validation set was used for hyperparameter tuning. The results below correspond to the performance of MARSY on the test set. Best performance values are in bold-face and underlined. The rows are sorted based on PCC of $S_{mean}$.

| Model | Cancer Type | Training Set Size | ZIP | | | $S_{mean}$ | | |
|---|---|---|---|---|---|---|---|---|
| | | | SCC | PCC | RMSE | SCC | PCC | RMSE |
| MARSY (MCF7) | Breast Cancer | 81,294 | 0.688 | **0.845** | **5.36** | **0.816** | **0.864** | **7.45** |
| MARSY (MFP) | NA | 154,718 | 0.699 | 0.834 | 5.46 | 0.812 | 0.858 | 7.65 |
| MARSY (PC3) | Prostate Cancer | 79,588 | **0.704** | 0.836 | 5.48 | 0.794 | 0.853 | 7.73 |
| MARSY (A549) | Lung Cancer | 55,610 | 0.680 | 0.838 | 5.46 | 0.792 | 0.852 | 7.76 |
| MARSY (A375) | Skin Cancer | 1,558 | 0.140 | 0.094 | 13.26 | 0.303 | 0.329 | 15.95 |

**Table S5:** The performance of MARSY against three other state-of-the-art models using Morgan fingerprints (MFP). The same validation and test sets used for results reported in Table S4 are used here. The validation set was used for hyperparameter tuning of all models. The results below correspond to the performance on the test set. Best performance values are in bold-face and underlined.

| Model | Drug Features | Training Set Size | ZIP | | | $S_{mean}$ | | |
|---|---|---|---|---|---|---|---|---|
| | | | SCC | PCC | RMSE | SCC | PCC | RMSE |
| MARSY | MFP | 154,718 | **<u>0. 699</u>** | **<u>0.834</u>** | 5.46 | **<u>0.812</u>** | **<u>0.858</u>** | 7.65 |
| MatchMaker | MFP | 154,718 | 0.628 | 0.821 | **<u>5.43</u>** | 0.787 | 0.854 | **<u>7.60</u>** |
| TreeCombo | MFP | 154,718 | 0.591 | 0.772 | 6.08 | 0.750 | 0.835 | 8.00 |
| DeepSynergy | MFP | 154,718 | 0.526 | 0.777 | 6.26 | 0.766 | 0.838 | 8.07 |

**Table S6:** ZIP scores of 4692 drug-pairs for 75 cancer cell lines, provided as a separate xlsx file. Both the training data and predicted scores are provided as different tabs.

**Table S7:** The difference between ZIP scores of breast cancer CCLs versus other CCLs based on the imputed synergy scores of 4692 drug-pairs for 75 cancer cell lines, provided as a separate xlsx file.
